# Evolutionary genomics reveals plant-associated origins of acetic acid bacteria in fermented food

**DOI:** 10.1101/2025.07.09.663944

**Authors:** Ana Cuesta-Maté, Thibault P. C. Bret, David A. Duchêne, Angela M. Oliverio, Jonathan Z. Shik, Sandra B. Andersen, Robert R. Dunn, Juan Antonio Rodríguez, Veronica M. Sinotte

**Author notes:** Equal contribution (Juan Antonio Rodríguez, Veronica M. Sinotte).

## Abstract

Humans have historically relied on acetic acid bacteria (AAB) for food fermentation, yet their origins likely trace back to free-living species outside of human environments. In nature, plants, fruit flies, and social insects host AAB. However, the evolutionary transitions of AAB from symbiotic hosts to fermented foods remain ambiguous. Here, we conduct a comprehensive phylogenomic analysis of 570 publicly available AAB genomes. Unlike other ferment-associated microbes, ferment-associated AAB appear to have exclusively evolved from clades inhabiting plants, and never social insect hosts. In particular, the sugar-rich fruits and flowers of angiosperms appear to promote diversification of AAB and transitions to fermented food habitats. Genomic features are similar in plant- and ferment-associated AAB; conversely, social insect-associated AAB have reduced genome sizes and number of genes, which may have limited their functional capability to disperse into ferments. Plant- and ferment-associated AAB share in the ability to metabolise diverse plant carbohydrates, though both produce habitat-specific carbohydrate-active enzymes. In contrast, metabolic capacity is reduced in social insect-associated AAB. By tracing the phylogenomics of this clade, we understand how evolution forged AAB capable of performing metabolic work for humans, shaping the past and potential futures of fermentation.

## Introduction

Acetic acid bacteria (AAB; Acetobacteraceae, acetous group) are central to the fermentation of food and drinks (Guzman & Vilcinskas, 2019). Human populations have harnessed the potential of AAB for food fermentation over the last several thousand years (Mazza & Murooka, 2009; Giudici et al., 2015a; Pecci et al., 2020). In the history of Western science, AAB were among the first microbes characterised; they were documented by Antonie van Leeuwenhoek in his studies of vinegar (Lane, 2015; van Leuwenhoek, 1676). Today, AAB are fundamental to fermentations including vinegar, kombucha (De Roos & De Vuyst, 2018; Tran et al., 2020), select sourdoughs (Landis et al., 2021; Rappaport et al., 2024), and vitamin C biosynthesis (Xu et al., 2004; Hattori et al., 2012; Gomes et al., 2018; La China et al., 2018). In ferments, these bacteria largely metabolise sugar and alcohol into acetic acid. This acid reduces substrate pH, inhibits the growth of other microbes, and thus facilitates food preservation (Matsushita & Matsutani, 2016; Rahman, 2020). However, before human use, AAB evolved as symbionts of plants, social insects (i.e., bees and some ants), and non-social insects (i.e., fruit flies) (Yamada & Yukphan, 2008; Chouaia et al., 2014; Pedraza, 2016; Brown & Wernegreen, 2019; Härer et al., 2022; Miller, 2023; Sannino & Dobson, 2023).

Many of the microbes used by humans in fermentation have been domesticated and controlled through various longstanding culinary practices. For example, microbes for fermentation are added via microbial starters (i.e. commercial bacterial cultures) (De Melo Pereira et al., 2020), previous ferments (i.e. backslopping) (Landis et al., 2021) or the repeated use of tools (i.e. barrels) (Verberg, 2019; Reese et al., 2020; Dunn et al., 2021) or spaces (i.e. fermentation rooms, wine/cheese caves) (Bokulich & Mills, 2013). Prior to the advent of food fermentation by humans, these microbes lived and evolved outside of human environments. The pre-human evolutionary origins of microbes used to produce fermented foods have started to become documented. For example, the brewer’s yeast *Saccharomyces cerevisiae* depends upon social wasps for dispersal, overwintering, and hybridization (Stefanini et al., 2012, 2016; Madden et al., 2018, 2022). Stefanini et al., (2012) suggest that such wasps were the original habitat for these yeasts. Today, wasps can still disperse yeast into natural wine fermentations, contributing to the “terroir” of the wine production (Di Paola et al., 2023). Similarly, ants have been suggested to be the original and natural habitat of the sourdough lactic acid bacteria, *Fructilactobacillus sanfranciscensis* (Sinotte et al., 2025; Zheng et al., 2022). It is possible that AAB also evolved in association with insect or plant hosts and were then co-opted from these habitats by humans in the context of fermentation.

We hypothesise that coevolutionary histories of AAB with symbiotic hosts provided adaptations that predisposed a subset of AAB to successfully disperse into fermented foods. Ancestral symbioses of AAB with social insects that feed on sugar-rich resources (Anderson et al., 2013, 2018; Brown & Wernegreen, 2016, 2019; Smith & Newton, 2020) or with plants that produce sugary fruits or floral nectar (Pedraza, 2008, 2016) could have prepared AAB to overtake sugar-rich fermented food environments. Fruit flies may have also mediated this AAB transition into fermented foods because they host AAB and frequently visit flowers, fruits, and ferments (Clymans et al., 2019; Madden et al., 2022; Woodcock et al., 2014). To date, studies of AAB have lacked sufficient interdisciplinarity to connect the dots between the applied uses of AAB in food fermentation (Guzman & Vilcinskas, 2022; Murooka, 2016) and the natural history of their insect-(Anderson et al., 2013; Miller et al., 2021) and plant-associated AAB ancestors (Pedraza, 2016), siloed within these disciplines.

We hypothesise that evolutionary transitions of AAB from symbiotic hosts to fermented foods required genomic and metabolic adaptations. We predict that genomic signatures of adaptation to consistent environments and microbial inheritance observed in microbe-host associations extend to AAB associated with fermented foods. We further predict that metabolic adaptations, specifically related to carbohydrates, may occur in the AAB habitat transition to fermented foods. For example, the difference in the carbohydrate landscape between floral nectaries to sourdough (Lievens et al., 2015; Gobetti et al., 2019; Nicolson, 2022) may prompt metabolic evolution. Thus, genomic signatures in AAB found in ferments may include 1) genome reduction (Moran et al., 2008), 2) AT-nucleotide bias (Van Leuven & McCutcheon, 2012; Brown & Wernegreen, 2016), 3) reduced gene number or functionality (i.e. pseudogenes) (Giovanoni et al., 2014; Moran & Bennet, 2014), and thus 4) metabolic adaptations.

Here, we use comparative phylogenomics of AAB associated with ferments, plants, social insects and fruit flies to clarify their evolutionary history and ancestral ecological associations. First, we reconstruct an AAB phylogeny to infer the evolutionary origins of AAB used in fermented foods. Second, we test whether AAB transitions from symbiotic hosts to fermented foods yield evolutionary signatures of 1) genome reduction, 2) AT-bias, and 3) reduced gene number and functionality. Third, we compare the metabolic profiles of AAB, focusing on carbohydrate metabolism, to investigate whether AAB show 4) changes in their metabolic ability reconcilable with adaptation to life within ferments. Ultimately, this study underscores the relevance of evolutionary genomics, which provides a lens through which to discern the origins of fermented food AAB and, importantly, clarify how these diverse origins may be implicated in past and future food fermentations.

## Results

### Fermented food AAB originate from plant symbionts

Phylogenetic inference based on acetic acid bacteria (AAB) genomes reveals that ferment-associated bacteria have exclusively evolved from clades inhabiting plants, and not from clades found in insects (Figure 1A, Supplementary Figure 1). All AAB associated with fermented foods are nested within one of four lineages also associated with plants: *Komagataeibacter*, *Acetobacter*, *Gluconobacter,* and *Asaia*. Our phylogeny suggests that transitions from plant to fermented food environments occurred through both recent dispersal events and from ancient plant-associated clades that were, in some way, predisposed to success in ferments. For example, *Komagataeibacter*, a lineage found with both ferments and plants has an initial divergence from *Gluconoacetobacter*, a strictly plant-associated lineage, suggesting that some attribute of *Komagataeibacter*’s ancestor but not of *Gluconoacetobacter*’s facilitated fermentation. The remaining plant-associated AAB clades then appear to be primed for dispersal to fermented food habitats. We find two social insect-associated AAB lineages: *Commensalibacter* and *Bombella*. The *Commensalibacter* lineage has primarily been detected in social insects; a handful of strains have also been isolated from other insects, flowers, and berries, perhaps suggesting dispersal from bees visiting floral nectaries and fruits/berries (Botero et al., 2023; Guzman et al., 2021; Cuesta-Maté et al., in prep). Species of the *Bombella* clade are, at least based on sampling to date, confined to the social Hymenoptera (Härer et al., 2022). We find no evidence that any strains from these lineages have colonised ferments.

**Figure 1.**
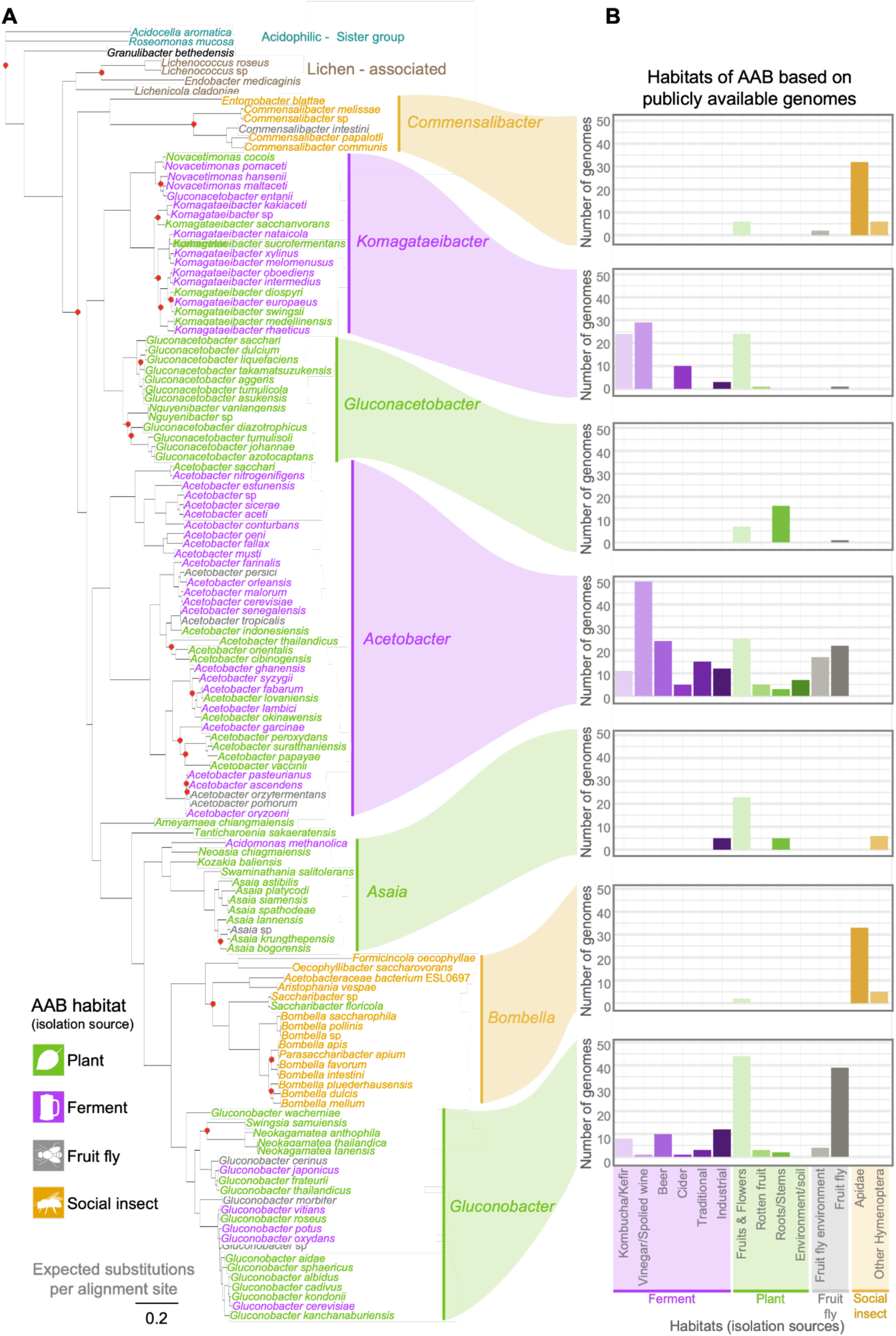
Phylogenomic tree of acetic acid bacteria (AAB) and their habitats. **(A)** The AAB phylogeny inferred using 134 genomes, with each genome being the most complete and least contaminated for its species (Supplementary Data 1). The tree was built on a 71 single-copy core gene alignment; nearly all branches displayed satisfactory site concordance factors (sCF) (>70, considered strong support; red dots identify the branches with <70sCF). The font colour of a species indicates the isolation source for the specific genome. Clade classification is according to Guzman & Vilcinskas, 2022. The shading of clades is based on the habitat from which the majority of the respective genomes have been isolated. **(B)** Barplots of the habitats are based on isolation sources of 570 publicly available genomes used in this study (Supplementary Data 2). Habitats are further distinguished by microhabitats of AAB isolates. For example, fermented food microhabitats include kombucha/kefir, vinegar/spoiled wine, beer, cider, traditional and industrial ferments. In the case of insect-associated AAB, social Hymenoptera (ants, bees and wasps) are separated from Diptera (fruit flies) due to the distinct evolutionary history of AAB and social insects. Within Hymenoptera, the main microhabitat is Apidae (bees).

An individual plant can contain many microhabitats. We hypothesised that it was specifically the dependence of AAB on the subset of plant microhabitats that are sugar-rich, such as fruits or floral nectaries, that predisposed AAB lineages to successfully colonise the similar sugary fermented foods. For example, floral nectaries are rich in simple sugars as sucrose, glucose, fructose, and potential yeast byproducts, and vinegar derived from wine also contains glucose, fructose, and ethanol from yeasts; thus the similarity between carbohydrate landscapes may allow AAB to disperse from sugar-rich plant microhabitats to ferments. To test this, we classified 570 AAB genomes deposited on NCBI according to their original isolation source and hence microhabitat. The plant-associated microhabitats of AAB can be divided into fruits and flowers, rotten fruits, roots and stems, or soil; ferment habitats of AAB, meanwhile, include vinegar, beer, kombucha, traditional ferments (i.e. agua de coco), or industrial ferments (i.e. biorefineries) (Figure 1B, legend). Consistent with our hypothesis, in all clades associated with ferments, the majority of plant-associated AAB were isolated from fruits and flowers (Figure 1B). Notably, in the only clade strictly associated with plants, *Gluconoacetobacter*, AAB are more often found in roots and stems than fruits and flowers, underlining that plant microhabitat likely dictates dispersal to ferments. In clades found in fruits, flowers, and ferments, AAB were also frequently associated with fruit flies, in keeping with the hypothesis that flies mediate dispersal by frequenting these microhabitats (Clymans et al., 2019; Madden et al., 2022; Woodcock et al., 2014). Overall, life in sugar- and ethanol-rich plant habitats, such as flowers, fruits, and rotting fruit, appears to have predisposed AAB to disperse to and be domesticated in fermentation habitats.s

We next estimated the evolutionary histories of AAB and symbiotic hosts, given that AAB appear to have evolved with angiosperms (flowering plants). Our AAB phylogeny was time-calibrated based on the divergence (80 Mya) between honeybees and stingless bees (Cardinal & Danforth, 2011), which appear to host distinct AAB lineages (Sarton-Lohéac et al., 2023; Mazel et al., 2025). This allows rough comparison of natural history events but not robust dating due to variable substitution rates and horizontal gene transfer in bacteria. The evolutionary history of AAB appears to be concordant with that of hosts and sugar-rich habitats (Supplementary Figure 2). AAB diversification aligns with the angiosperm terrestrial revolution, and the consequential accelerated ability of environmental sugars (Morreau et al 2006; Peris & Condamine, 2024). For example, the topology of the AAB tree shows the earliest split to occur with the lichen-associated genera (Noh et al., 2020; Pankratov et al., 2020) and divergence of the social insect-associated *Bombella* clade during corbiculate bee diversification (Cardinal & Danforth, 2013).

### AAB have genomic signatures of adaptation to consistent environments

Bacteria associated with consistent environmental conditions often adapt via reduction in genome size and GC nucleotide percentage. We predicted that such adaptations occur across AAB transitions to fermented foods because a) fermentations occur under controlled, consistent conditions (Dunn et al., 2021), b) fermentation often favours fast growth over metabolic plasticity, and c) fermentation practices (i.e. backslopping, repeated tool use) can encourage microbial inheritance (Bokulich & Mills, 2013; Verberg, 2019). Similar conditions within symbiotic hosts is known to prompt genomic evolution, albeit over longer evolutionary times (Van Leuven & McCutcheon, 2012; Giovanoni et al., 2014; Moran & Bennet, 2014; Brown & Wernegreen, 2016). Contrary to our predictions, we did not observe overall differences in genome size or GC content in AAB found in ferments compared to their plant origins (Figure 2A).

**Figure 2.**
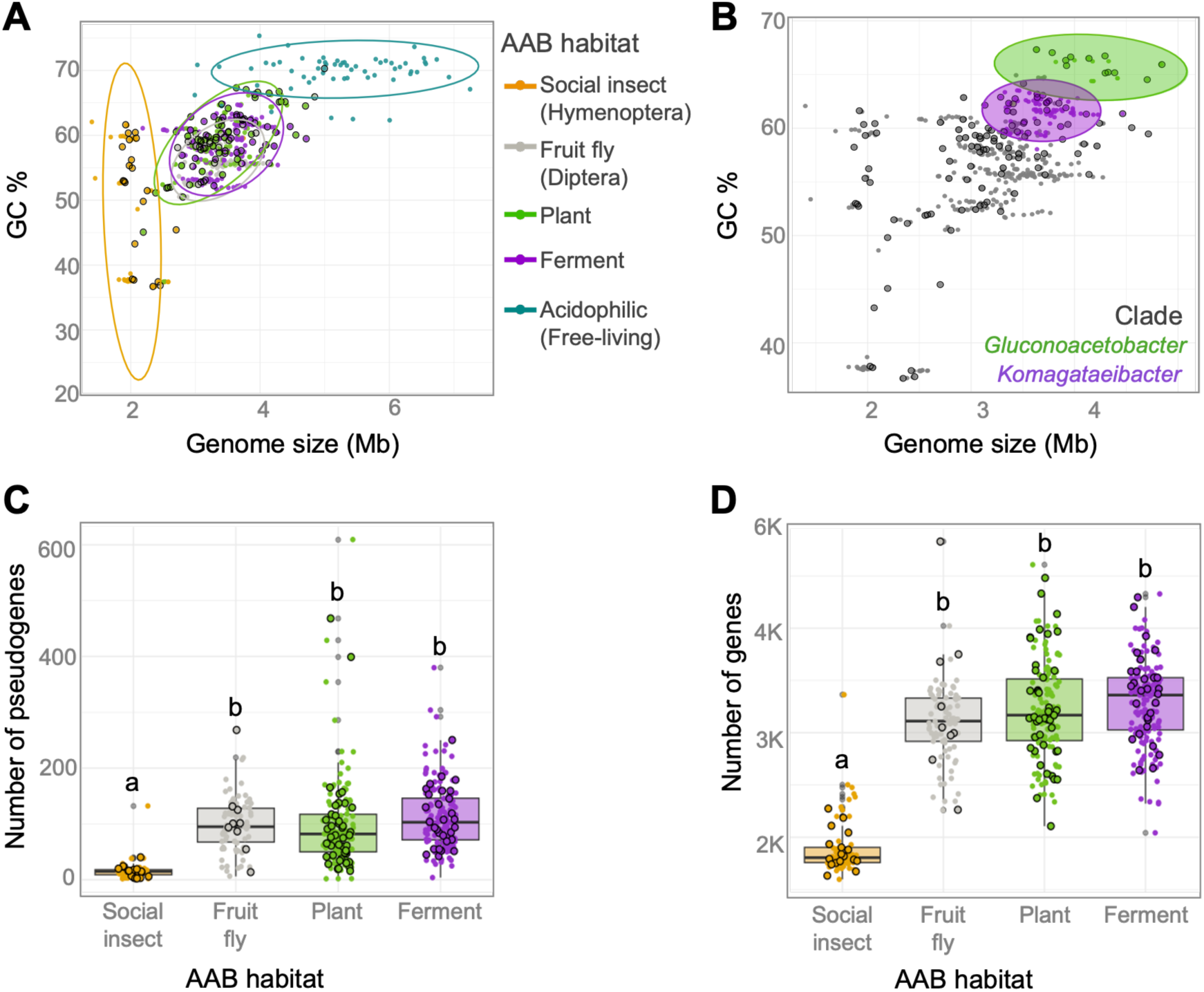
Genomic characteristics of acetic acid bacteria found in fermented foods, plants, fruit flies and social insects. (**A**) Genome size and GC% of 570 publicly available AAB genomes, and 87 non-AAB acidophilic bacteria genomes (Supplementary Data 2). Data points with the black border represent the 132 genomes included in the tree (Figure 1); for those, we tested the heritability of the genomic traits here studied, by performing Blomberg’s K test. This test quantifies the phylogenetic signal K; K>1 the trait is more similar than expected under brownian motion (preferentially selected), K<1, the trait is less similar than expected under brownian motion (preferentially lost). Phylogenetic signal is significant for both genome size (Blomberg’s *K*=0.8684; *p*<0.001) and GC content (Blomberg’s *K*=1.9264; *p*<0.001). For those 132 genomes, AAB habitat also has a significant effect on genome size (phylogenetic generalised least square analyses (PGLS) λ=0.947, *p*=0.0015); **(B)** Comparison between *Gluconacetobacter* (plant) and *Komagataeibacter* (ferment) sister clades, indicating reduction in genome size and GC% in the ferment clade. Grey dots represent the remaining AAB genomes not included in the comparison. **(C)** Total number of pseudogenes within AAB genomes. The phylogenetic signal is significant for the number of pseudogenes per genome (Blomberg’s *K*=0.1049, *p*=0.008). **(D)** Number of genes per genome. The phylogenetic signal is significant for the number of total genes per genome (Blomberg’s *K*=0.2598, *p*=0.001). The outlier, *Komagataeibacter kakiaceti,* contained 1094 pseudogenes and was removed. In panels **(C-D)** boxplots represent the first and third quartiles, the horizontal line represents the median, whiskers extend 1.5 interquartile ranges, and dots represent each of the data points. The letters represent statistical differences among groups according to PGLS with a Tukey Post Hoc test for pairwise comparisons (*p*<0.05). If the boxplots share the same letters, no statistical differences were found between groups. All statistics included in Supplementary Table 1.

We did, however, observe cases of genomic adaptation in AAB in relation to symbiotic transitions. We compared the genome size and nucleotide composition of AAB, found in symbiotic or fermented food habitats, to their sister clade of acidophilic bacteria, found free-living in soil and water. In the transition from free-living habitats, AAB reduced in genome size and GC content. Subsequent AAB evolution within these habitats significantly affected genome size (Figure 2A; Phylogenetic generalised least square analyses (PGLS) λ=0.947, *p*=0.0015) and non-significantly in GC content (Supplemental Figure 3A, 3B), suggesting genomic adaptation to more consistent symbiotic lifestyles, which has been previously observed (Figure 2A; Guzman & Vilcinskas, 2021). We then compared plant-associated AAB in the *Gluconacetobacter* clade, found largely in roots and stems, to the *Komagaetibacter* clade, found largely in fruits, flowers, and ferments. We observed that the *Komagataeibacter* clade has undergone a local weak reduction in genome size (Figure 2B, Supplemental Figure 3C; PGLS, λ:0.4134, *p*=0.0595), potentially driven by the clear reduction in the number of genes in *Komagataeibacter* (Supplemental Figure 3D; PGLS, λ:0.0000, *p*=0.0011), and a reduction in GC content (Supplemental Figure 3E; PGLS, λ:0.0000, *p*<0.0001). This reduction in *Komagataeibacter* may be driven by two differences in these habitats. First, fruits, flowers, and ferments may provide more consistent or narrow environmental conditions, for example in carbohydrate profiles. Second, in these sugar-rich environments, fast growth may be favoured over metabolic plasticity.

The most drastic and significant genomic changes were observed in social-insect associated lineages. Clades *Commensalibacter* and *Bombella* exhibit significant reductions in genome size compared to ferment and fly-associated AAB (Figure 2A, Supplementary Figure 3B, Supplementary Table 1; PGLS, all comparisons *p*<0.05). This appears to be due to the loss of both pseudogenes (Figure 2C; PGLS λ:0.2224, *p*<0.0001) and functional genes (Figure 2D; PGLS λ:0.8773, *p*<0.01), suggesting high genome density of social insect-associated AAB. Overall, the observed genomic adaptations in social insect-associated AAB are consistent with previous findings in the literature (Brown & Wernegreen, 2016, 2019; Alonso et al., 2019; Guzman et al., 2021; Härer et al., 2022; Botero et al., 2023); it appears that once AAB adapt to life within social insects, transitions to other habitats are unlikely to occur.

### Carbohydrate metabolism evolves with AAB transitions to fermented foods and social insects

We predicted that AAB transitions from symbiotic hosts to fermented food will have impacted carbohydrate metabolism. The carbohydrate profiles in fermented foods may differ from ancestral states, and it is the degradation of carbohydrates and consequent acid production that allows AAB to outcompete other microbes in their natural environments. AAB transitions to fermented food and social insect habitats appear to have modified the diversity of enzymes for carbohydrate metabolism, albeit in different ways. Ferment-associated AAB maintain similar numbers of carbohydrate-active enzymes (CAZymes) to those in their plant-associated counterparts (Figure 3A), but their composition differs (Figure 3B; Adonis pairwise plant vs ferment: F=3.1677, R^2^=0.0307, *p*=0.03). The differences in CAZyme composition are due, in part, to habitat-specific CAZyme families; 18 CAZyme families were unique to plant-associated AAB and six are unique to ferment-associated AAB (Supplementary Figure 4). Conversely, social insect-associated AAB carbohydrate metabolism appears specialised, with a reduction of CAZymes compared to plant or ferment-associated AAB, that becomes non-significant once we account for phylogenetic relatedness (Figure 3A; PGLS, λ:0.9333, *p*<0.09), and distinct CAZyme profiles from all other AAB (Figure 3B; Adonis pairwise, social insect vs all other habitats: *p*<0.01; Supplementary Table 2). As a result, plant- and ferment-associated AAB can putatively degrade a broad range of carbohydrate substrates, while this is limited in social insects (Figure 3C).

**Figure 3.**
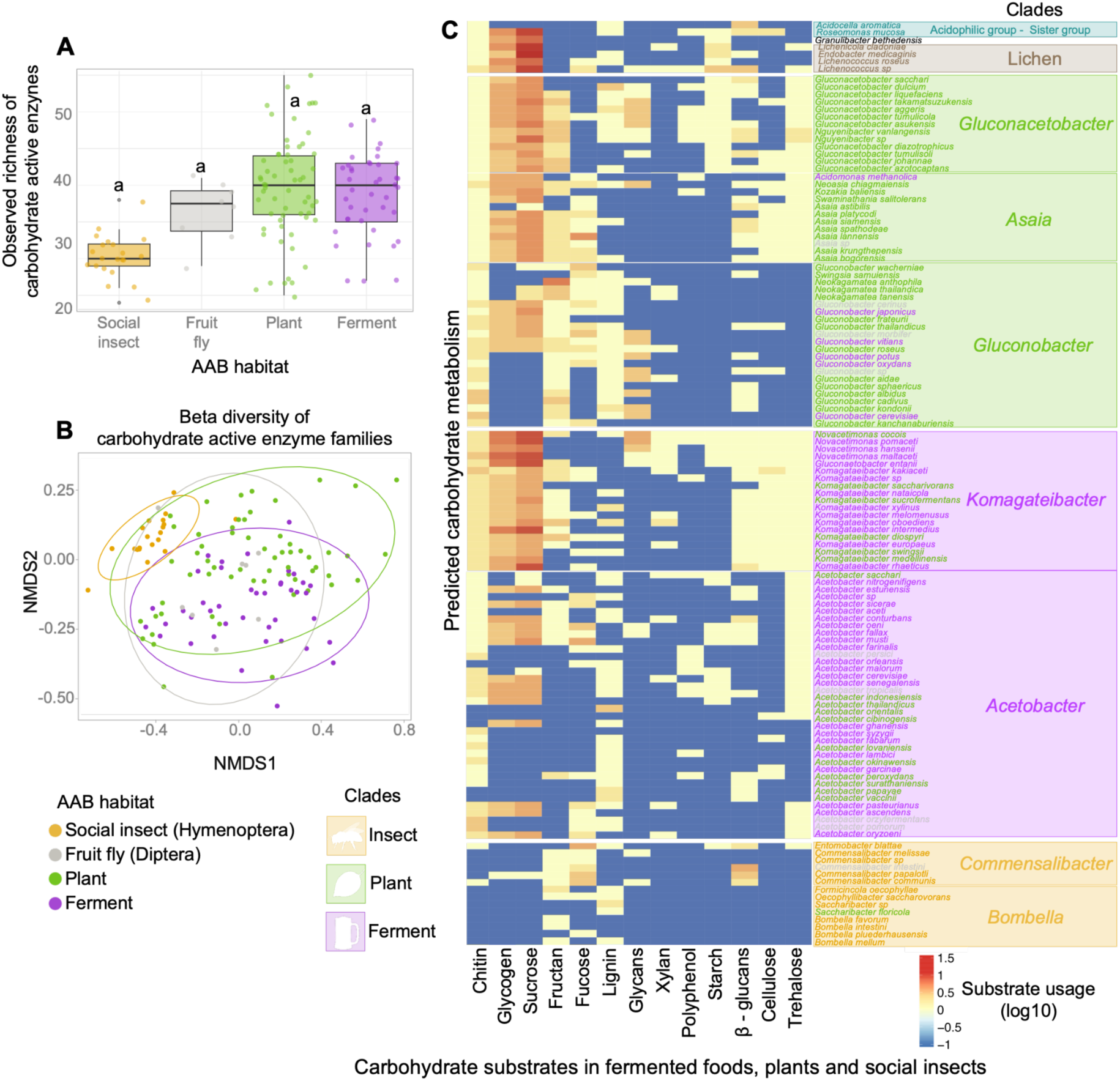
Carbohydrate metabolic potential of acetic acid bacteria isolated from fermented foods, plants, and insects. The 134 genomes used to reconstruct the AAB phylogenetic tree (Figure 1) were annotated for the study of metabolic capacity. **(A)** Observed richness of carbohydrate-active enzyme (CAZyme) families for each AAB habitat, represented as the absolute number of CAZymes annotated for each genome. 112 CAZymes were found across all genomes (Supplemental Data 3). Phylogenetic signal is significant for the absolute number of CAZymes encoded in each genome (Blomberg’s *K*=0.8598 *p*=0.001), though AAB habitat did not affect the total number of CAZymes PGLS λ=0.933, *p*=0.086). Boxplots represent the first and third quartiles, the horizontal line represents the median, whiskers extend 1.5 interquartile ranges and dots represent each of the data points. The letters represent statistical dissimilarities across groups according to Tukey Post Hoc pairwise comparisons (*p*<0.0001). All statistics are included in Supplementary Table 2. **(B)** Non-metric multidimensional scaling (NMDS) ordination based on Bray-Curtis distances of the CAZyme family profiles of each bacterial genome from the respective AAB habitats. The composition of CAZyme families also differed as a function of AAB habitat (Figure 3B; PERMANOVA: F_3,127_=10.209, R2=0.194, *p*<0.0001). **(C)** Heatmap of the number of CAZymes predicted to metabolise carbohydrate substrates found in fermented foods, plants and social insects.

## Discussion

In the context of fermentation, acetic acid bacteria (AAB) are among the most economically and culturally important microorganisms to humans. We show evidence that AAB in fermented foods evolved exclusively from AAB living with and on plants, in particular, angiosperms. We hypothesise that the similarities between ferments and sugar-rich plant parts facilitated these habitat transitions. First, many ferments are made from plants and can contain similar carbohydrates, perhaps explaining the overlap in carbohydrate metabolism. Second, ferments, floral nectar, and fruits host comparable microbiomes - both host AAB that engage in metabolic exchanges with yeast (De Roos & De Vuyst, 2018; Álvarez-Pérez et al., 2019; De Vuyst & Leroy, 2020; Fleet, 2003; Ge et al., 2025; Hall et al., 2018; Landis et al., 2021). This is in line with our observation that plant- and ferment-associated AAB have the ability to metabolise chitin and glycogen, carbohydrates found in yeast and other fungi. Third, AAB evolved to compete for ephemeral sugar-rich resources, which may occur in both angiosperms and fermented foods. Ephemerality is known to drive extreme resource competition (Smid & Lacroix, 2013; Bauer et al., 2018; Toju et al., 2018; Wittman & Fukami, 2018) and foster a “sit-and-wait” strategy. During substrate sc metabolic state (Walther & Ewald, 2004; Sundberg et al., 2014; Rayner et al., 2024). When resources become available, these microbes diarcity, AAB potentially wait in a lowvide quickly and produce acetic acid as a biocide (Matsushita & Matsutani, 2016; Tran et al., 2020; Wang et al., 2024; Han et al., 2024). Thus AAB may have initially evolved to monopolise seasonally flowering and fruiting angiosperm environments, making them also competitive in ephemeral human-produced ferments.

Traditional balsamic vinegar provides an excellent example of how similarities between plants and ferments may have facilitated AAB habitat transitions. The AAB species used in traditional balsamic vinegar fermentation belong to the same genera of AAB found on the grapes from which it is made (i.e. *Komagataeibacter*, *Novacetimonas*, *Acetobacter*) (Gullo et al., 2006; Gullo & Giudici, 2008; Murooka, 2016). Plausibly, the ancestral plant-associated AAB in those grapes bacteria were able to metabolise the environmental sugars and evolved in concert with *Zygosaccharomyces* yeasts, found on both grapes and in balsamic fermentation (Giudici et al., 2015b; Rainieri & Zambonelli, 2009; Solieri & Giudici, 2009; Mezzasalma et al., 2017). For example, in grapes or vinegar the AAB may metabolise ethanol produced by the yeasts and, based on our results, potentially have the ability to breakdown chitin and glycogen in the yeasts themselves. We hypothesise that a “sit-and-wait” strategy to quickly monopolise rotting grapes represented a pre-adaptation that enabled those first AAB to dominate when transferred to balsamic fermentation. The time of the first persistent habitat transition, from grapes to balsamic, is not known. However, centuries of use of the same wooden casks in traditional Italian balsamic production (Giudici et al., 2015) likely shaped the AAB evolution into the proficient balsamic AAB embedded within them. Thus, this is an exemplary case of how the microbial ecology of sugar-rich fruit environments affects the habitat transitions and evolution of AAB in fermentation habitats.

In AAB, we see modest genomic changes in some ferment lineages and dramatic genomic modifications in social insect lineages, aligned with adaptation to consistent environments, acidity, and in some cases vertical transmission. Both ferments and social insects can provide consistent environmental conditions for AAB, but tens of millions of years of social insect-AAB symbiosis appear to have driven a more drastic genome evolution. The low pH in fermented food and social insect environments may have also affected genome structure. Genome streamlining has also been recently suggested as a potential adaptation to life at low pH (Cortez et al., 2022). While ferment-associated AAB have only recently undergone their transition into lower pH environments, social insect host environments are typically acidic. For example, the clade *Bombella* may be found in bee honey or formicine ant crops, which can have pH ranges of pH 3.2-4.5 (Bogdanov et al., 2008) and pH 2-5 (Tragust et al., 2020), respectively.

We consistently find that AAB associated with fruit flies (*Drosophila*) are phylogenetically, genomically and metabolically similar to plant- and ferment-associated AAB. These similarities suggest that fruit flies exchange and transmit AAB between them, but they are unlikely permanent hosts due to their unstable microbiome composition (Broderick & Lemaitre, 2012). *Drosophila* flies are attracted to the natural sugars and yeasts found in plants and ferments. Natural sugar sources can include floral nectar (Woodcock et al., 2014), overripe or fermented fruits (Zhu et al., 2003; Kim et al., 2023), and human-made fermented products (Kleiber et al., 2014; Cha et al., 2012). Yeast found in sugary plant parts (i.e. ripe fruit) and ferments produce volatile compounds to attract fruit flies and use them as a dispersal vector (Palanca et al., 2013; Christiaens et al., 2014; Becher et al., 2018). Fruit flies may also be engaging in a similar diffuse mutualism with AAB, where the microbes’ honest signal of fermenting sugars is rewarded with transportation to new ephemeral sugar sources (Madden et al., 2018). The absence of fruit fly-specific carbohydrate enzymes, despite the accumulation of ferment-specific carbohydrate enzymes, further supports the instability of these associations, and implies metabolic changes similar to those observed in habitat transitions to fermented foods (Somerville et al., 2024).

AAB in social insects do not disperse to ferments, contrary to our predictions based on observations from other fermented food microbes (Stefanini et al., 2012, 2016; Di Paola et al., 2023; Sinotte et al., 2025). Rather, evolution in social insects has finely attuned AAB to their hosts. The apparent transition of AAB from plants into social insects occurred twice, in the clades *Commensalibacter* and *Bombella*. Evolution within social insects led to significant gene loss and genome reduction, consistent with prior findings of adaptive genome reduction in insect symbionts (Chouaia et al., 2014; Moran & Bennett, 2014; Latorre & Manzano-Marín, 2017; Comandatore et al., 2021). Further, it seems that once associated with their insect hosts, AAB evolve several times faster than their plant- or ferment-associated counterparts, potentially evincing a quick and defining adaptation to these environments (Brown & Wernergreen, 2016). The remaining genes appear to complement the hosts’ needs, favouring cooperation and avoiding conflict (McCutcheon & Moran, 2012; Wernegreen, 2015; Fisher et al., 2017). For instance, social insect-associated AAB cannot metabolise sucrose, a sugar prevalent in the diet of bees (Wang et al., 2021); or chitin, a structural component of insects’ exoskeletons (Chaudhari et al., 2011). Once AAB adapted to life with their social insect host, we hypothesise that their diminished metabolic capabilities prevented transitions to any other habitats.

Understanding the origins and ecologies of fermentative AAB present exciting avenues for fermented food research. This analysis of publicly available AAB genomes reveals a previously unseen diversity that may support novel applications in fermentation. Our findings suggest that fruits and flowers are an overlooked and biodiverse reservoir of AAB. The broad metabolic potential encoded by these AAB offers an opportunity to select strains that can be implemented in the fermentation of novel foods. For example, food waste products (i.e. sidestreams), such as brewer’s spent grain, contain recalcitrant carbohydrates including lignin and hemicellulose (Mussatto et al., 2006). Here, we identify AAB species that encode the metabolic machinery to potentially act on these carbohydrates. These AAB may thus allow sidestreams to be integrated in fermentations such as sourdough bread (Rappaport et al., 2024), supporting a circular economy and a sustainable food system (Fassio & Tecco, 2019; Tamasiga et al., 2022; Zhang et al., 2022). Second, we predict that the genes or enzymes under positive selection during transitions to fermentation have valuable functions for the transformation of food substrates. It offers an opportunity to identify and implement such AAB strains or enzymes to accelerate microbial domestication or support precision fermentation.

Evolutionary ecology allows us to engage with AAB diversity to effectively engineer future ferments and biosolutions, while uncovering how their evolutionary histories have shaped our current foods. We demonstrate how transitions from sugar-rich plant environments gave rise to ferment-associated lineages, while long-term associations with social insects drove specialised genome evolution. This framework highlights plant-associated AAB as a reservoir of metabolic diversity with potential for future food fermentations and biotechnological applications. More broadly, integrating evolutionary ecology into fermentation research can guide the discovery and engineering of microbial functions for next-generation fermented foods and sustainable biosolutions.

## Data and code availability

All genomes were downloaded from the National Center for Biotechnology Information (NCBI) page, and the accession numbers are available in Data S1. This paper does not report any original code. All code used is available at github.com/ThBret/Fermentation-bacteria and will be published with a doi through Zenodo. All additional data generated is available in Supplemental Data are available on Mendeley data repository (https://data.mendeley.com/preview/gv45c4nn3f?a=36a84de3-e7ef-4b01-87c7-75a155e9f1c6; to be published upon acceptance).

## Supplemental information

● Supplemental Document. Figures S1-S4. Tables S1 and S2.
● Data S1. Genomes included in the phylogenomic tree.
● Data S2. Genomes included the genomic characteristic analysis.
● Data S3. Annotated CAZymes.

## Supporting information

Figure S2. Time-calibrated phylogenomic tree of acetic acid bacteria (AAB) and their habitats, related to Figure 1.

Figure S3. Pairwise comparisons of genomic characteristics within AAB habitats, related to Figure 2.

Figure S4. Carbohydrate active enzymes (CAZyme) potential of AAB across the phylogeny, related to Figure 3.

Figure S1. Ancestral state reconstruction of the habitat trait, related to Figure 1.

Table S1. Pairwise comparisons of genomic characteristics of AAB genomes from Social insects, Fruit fly, Plant and Ferments, related to Figure 2.

Table S2. Carbohydrate active enzymes (CAZyme) potential of AAB across from Social insects, Fruit fly, Plant and Ferments, related to Figure 3.

## Acknowledgments

The authors would like to thank Jazmín Ramos-Madrigal for her theoretical contributions and methodological discussions early in the development of this project. The authors thank Jennifer Wernegreen and Leonie Johanna Jahn for their valuable theoretical contributions in the earlier stages of the project. The authors thank Robert Murphy for his input on the analyses of CAZyme activity. The authors thank Dinah Maran Parker, Joshua Evans, and Margarita M. López-Uribe for comments on earlier drafts of the manuscript. A.C.M., T.P.B., S.B.A. V.M.S. was supported by the Danish National Research Foundation Centre for Evolutionary Hologenomics DNRF grant nr 143. J.A.R reports financial support was provided by H2020 Marie Curie-Sklodowska Actions PF award number 101064828 (POPARCH). D.A.D. acknowledges funding from a Novo Nordisk Fonden Emerging Data Science Investigator Award (NNF23OC0084647). A.M.O. was supported by National Science Foundation MCB award #2328528. J.Z.S. was funded by the Villum Foundation (VEX grant no. 50281), and a Semper Ardens: Accelerate grant (CF22-0664) from the Carlsberg Foundation, Denmark (https://www.carlsbergfondet.dk).

## Author contributions

A.C.M., R.R.D. and V.M.S. conceived the paper, A.C.M., T.P.B., D.D., J.A.R. and V.M.S. conceived the analyses, A.C.M. and T.P.B. collected the data, D.D., A.M.O., J.Z.S, S.B.A. contributed data or analyses tools, A.C.M., T.P.B. and D.D. performed the analyses. A.C.M., R.R.D, J.A.R. and V.M.S. wrote the paper, with continuous collaboration with the rest of the authors.

## Methods

### Genomic dataset and data curation

A dataset of the 675 publicly available genomes of *Acetobacteraceae* bacteria was curated for phylogenomic analysis. Genomes for all acetic acid bacteria, also known as the acetous group (Guzman & Vilcinskas, 2022; Qiu et al., 2021), were downloaded from NCBI Genbank in January 2024 (n=588). Additionally, the reference genomes of the acidophilic group (n=87), sister and basal to the AAB group and also within the *Acetobacteraceae* family, were downloaded. Genome size, guanine-cytosine content percentage (GC content), completeness, contamination scores, and the isolation source were retrieved for each of the genomes from the NCBI database. We supplemented the NCBI data with the bacterial temperature and pH ranges when available (REF: https://bacdive.dsmz.de/). Only genomes that were over 90% complete with <5% contamination (defined as the presence of foreign DNA sequences within a genome assembly, belonging to a different organism) were included in our study, with the exception of 7 genomes: *Acetobacter papayae*, *Acetobacteraceae bacteriumESL0697*, *Asia astibilis*, *Asia platycodi*, *Gluconacetobacter sacchari*, *Komagataeibacter kakiaceti*, *Nguyenibacter vanlangensis*) which were added for taxonomic completeness (Supplementary Data 2). In the end, 570 genomes were included for analysis of acetic acid bacteria genome characteristics.

### Phylogenomic analyses

The most complete and least contaminated genome for each of the species of AAB, under the criteria described above, was included for the construction of phylogenetic trees, leading to a total of 132 genomes. Two outgroups belonging to the basal acidophilic group were included, *Roseomonas mucosa* and *Acidocella aromatica* (Supplementary table 1). Anvi’o v.8 (Eren et al., 2021) was used to construct the *contigs.db* files, using the commands *anvi-script-reformat-fasta* and *anvi-gen-contigs-database*. A total of 71 single-copy core genes (SCG) were used for phylogenetic inference, as specified in the *Bacteria_71* Hidden Markov Model (HMM) profile (Hug et al., 2016; Lee, 2019) embedded in Anvi’o, using *anvi-run-hmms* over each *contigs.db* file. Amino acid sequences for SCG were obtained through *anvi-get-sequences-for-hmm-hits*, with the flag *get-aa-sequences*. The alignment was constructed from amino acid sequences and was composed of 18,698 columns including 8,661 parsimony-informative sites, *i.e.* columns that contained at least two different amino acids present in more than two sequences (Katoh et al., 2002; Ranwez et al., 2011). The alignment was used to construct a maximum-likelihood tree with IQ-Tree2 v.2.2.6 (Minh et al., 2025). Phylogenetic reconstruction was based on the best-performing LG+F+R5 substitution model, as determined using information-criterion model selection implemented in *ModelFinder* (Kalyaanamoorthy et al., 2017). To quantify support across branches we focused on the highly intuitive site concordance factors, which represent the portion of informative alignment sites that include an explicit signal supporting each given tree branch (sCF; (Minh et al., 2020). The package *ape* (Paradis et al., 2004) was used to visualise trees in R (v. 4.3.2).

### Trait evolution patterns

Phylogenetic signal refers to the phylogenetic clustering of a given trait, expected if the trait is highly heritable and arises rarely, as opposed to showing random scatter across the phylogenetic tree (Revell et al., 2008; Kamilar & Cooper, 2013). We investigated the phylogenetic signal of genome size, GC content, average gene length, proportion of coding sequences, number of genes and number of pseudogenes, using Blomberg’s *K* (Blomberg et al., 2003) as implemented in the *phylosig* function of the Phytools v2.1.1 R package (Revell, 2024). We then investigated the evolution of the isolation source trait by fitting a Markov-based model for the evolution of discrete traits (Lewis, 2001) using the *fitMk* function of the Phytools v2.1.1 R package (Revell, 2024). The function was run with an ARD model to obtain evolutionary rate values (*q*) for transitions between each isolation source.

### Time-dated phylogenetic inference

Phylogenetic time-calibration was based on one key split in the phylogeny, between *Parasaccharibacter apium* and *Acetobacteraceae bacterium ESL0697*, according to the divergence time of their hosts. *Parassaccharibacter apium* is isolated from honeybees *Apis mellifera*, while *Acetobacteraceae bacterium ESL0697* is isolated from stingless bees *Melipona lateralis*. Exchange of this bacterial symbiont between these two species is highly unlikely after their evolutionary divergence, given their high specificity to their insect host gut microbiomes and the lack of known interactions between distant geographic distributions (Powell et al., 2016; Kwong et al., 2017; Mazel et al., 2025). This time calibration is relaxed and distant enough that it seems plausible a divergence between these timepoints. We make the assumption that the timing of their divergence is also the time of the most recent common ancestor between their bacterial microbiomes. The split between these bee taxa is estimated to have occurred between 78 and 95 million years ago (Cardinal & Danforth, 2011), based on fossil records and estimates of the timing of the appearance of eusociality (Brady et al., 2006). Bayesian divergence time inference was performed by forcing the IQ-TREE2 tree topology, under approximate likelihood computation for highly efficient analysis as implemented in the module MCMCTree (dos Reis, 2022) of the PAML package v4.10.7 (Yang, 1997). We used a GTR+Γ model of nucleotide substitution and a relaxed clock with gamma-distributed rates across branches. A birth-death process was used as a tree prior with default parameters. We approximated the posterior distribution via Markov chain Monte Carlo sampling, using 10^7^ chain steps after a burn-in phase of 10^5^, with samples taken every 10^3^. Convergence to the stationary distribution was verified by running two independent analyses and comparing parameter estimates and checking that effective samples sizes across all parameters were above 200.

### Analysis of genomic characteristics

Genome size, GC content, completeness and contamination scores were also retrieved for each of the genomes. Further data curation was performed using Prokka v.1.14 (Seemann, 2014) for gene annotation and coding density. Boxplots and scatterplots were generated for several genomic features, and all statistical analyses were performed in R (v. 4.3.2). For the analyses of genomic characteristics such as genome size, GC content and coding sequence information, we realise that the phylogenetic relatedness of the genomes included in this study implies their lack of independence from each other. Thus, we account for the non-independence of our samples by performing a Phylogenetic Generalised Least Squares (PGLS) test. Following methods described by Angeli *et al*. (2025), we applied the *pgls* function of the R package *caper* (Orme et al., 2025) to assess relationships between genomic characteristics and ecological source while accounting for phylogenetic non-independence. The previously constructed phylogeny was incorporated into the input data so as to account for phylogenetic autocorrelation. Genomic traits evaluated included genome size, GC content, total gene count, number of pseudogenes and average gene length. Multicollinearity among predictors was assessed using the *vifcor* function of the R package *usdm* (Naimi et al., 2014), using the default correlation threshold value of 0.9. PGLS models were fitted to estimate Pagel’s λ values which were subsequently fixed in generalised least squares (GLS) models generated with *nlme* (Pinheiro et al., 2026). ANOVA tests and Tukey-adjusted pairwise comparisons were computed with *emmeans* (Lenth & Piaskowski, 2026).

### Carbohydrate metabolic potential of AAB

We explored the carbohydrate metabolism capabilities by using the dbcan4 v.4.0 pipeline (Zheng et al., 2023), which scanned our AAB genomes against a pre-annotated CAZyme nucleotide database (Drula et al., 2022). We calculated Observed richness and Shannon diversity on the relative abundance of CAZyme families in each of the AAB habitats, using the *estimateR* and *diversity* functions in the *vegan* R package (Oksanen et al., 2025). The effect of the isolation source was determined in R (v. 4.3.2) using the Phylogenetic Generalised Least Squares (PGLS) test, as described in the previous section. Compositional differences for CAZyme families between AAB genomes were determined by calculating Bray Curtis dissimilarity differences, and performing non-metric multidimensional scaling (NMDS) analysis, using the *metaMDS* function in the *vegan* package (v2.6-6.1, Oksanen, 2013) in R (distance=”bray”). The ordination of these results was visualised using *ggplot2* (v.3.5.1). To determine if the isolation source had an effect on AAB CAZyme profiles, we performed a Permutational Multivariate Analysis of Variance (PERMANOVA) using the *adonis2* function from *vegan* v2.6-6.1 on the distance matrix of Bray Curtis dissimilarity measurements. Pairwise comparisons of each timepoint were explored with the *pairwise.adonis2* function from the *pairwiseAdonis* package v.0.4.1. The heatmaps were reconstructed with the package pheatmap (Kolde, 2025) and *ComplexHeatmap* (Gu et al., 2015).

