## Supplementary figures and images for "Evolutionary genomics reveals plant-associated origins of acetic acid bacteria in fermented food"

### Figure S1. Ancestral state reconstruction of the habitat trait, related to Figure 1.

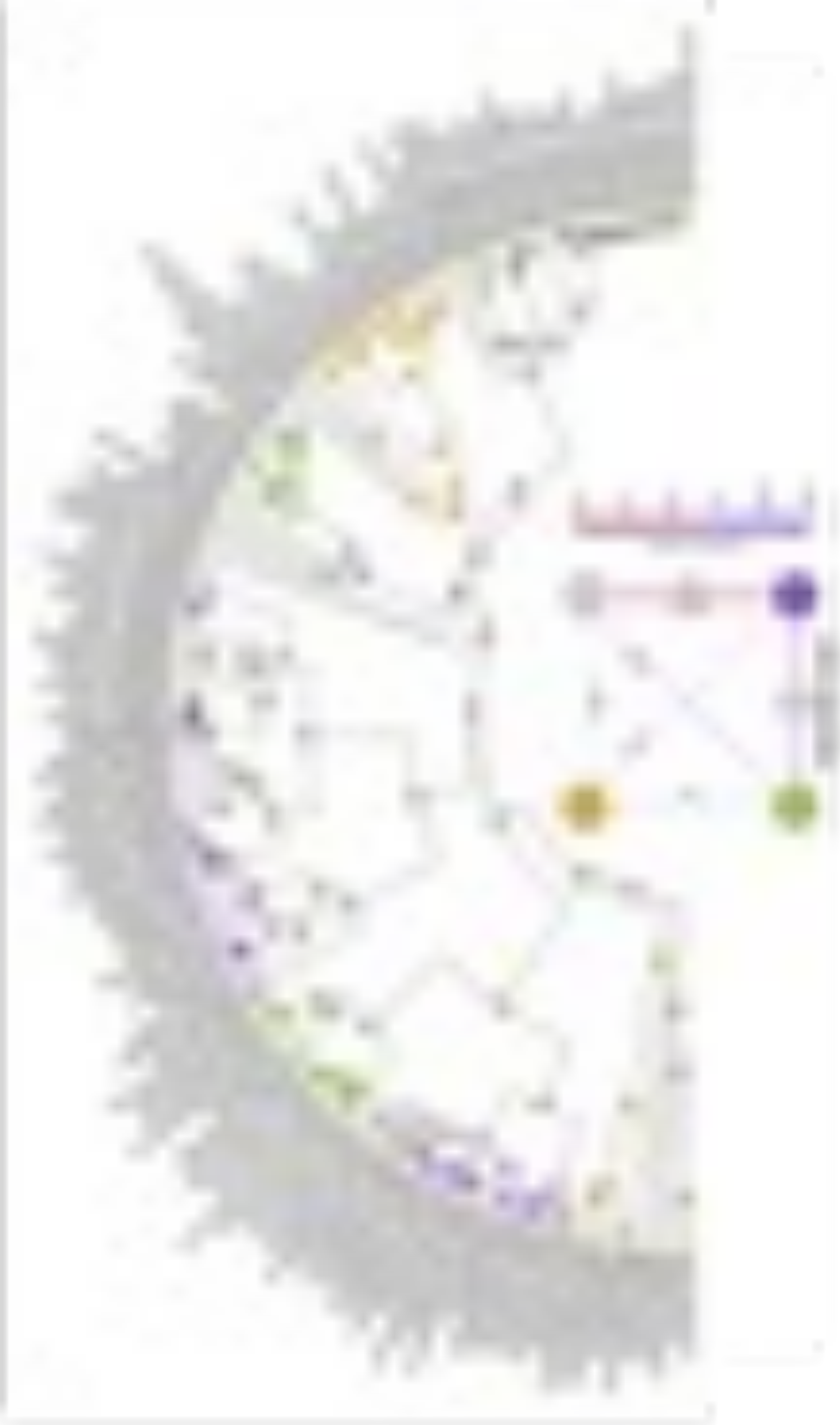

### Figure S2. Time-calibrated phylogenomic tree of acetic acid bacteria (AAB) and their habitats, related to Figure 1.

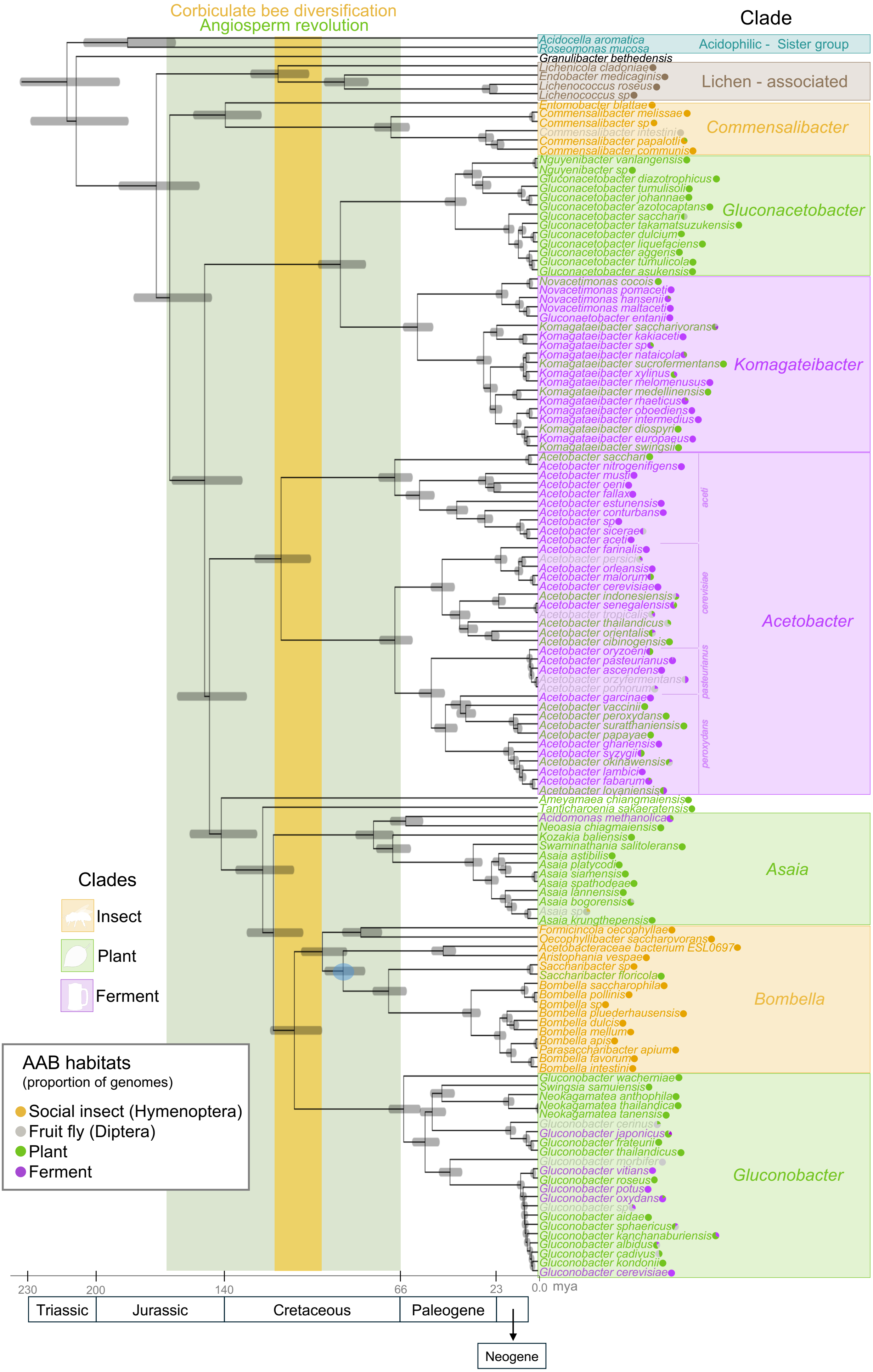

### Figure S3. Pairwise comparisons of genomic characteristics within AAB habitats, related to Figure 2.

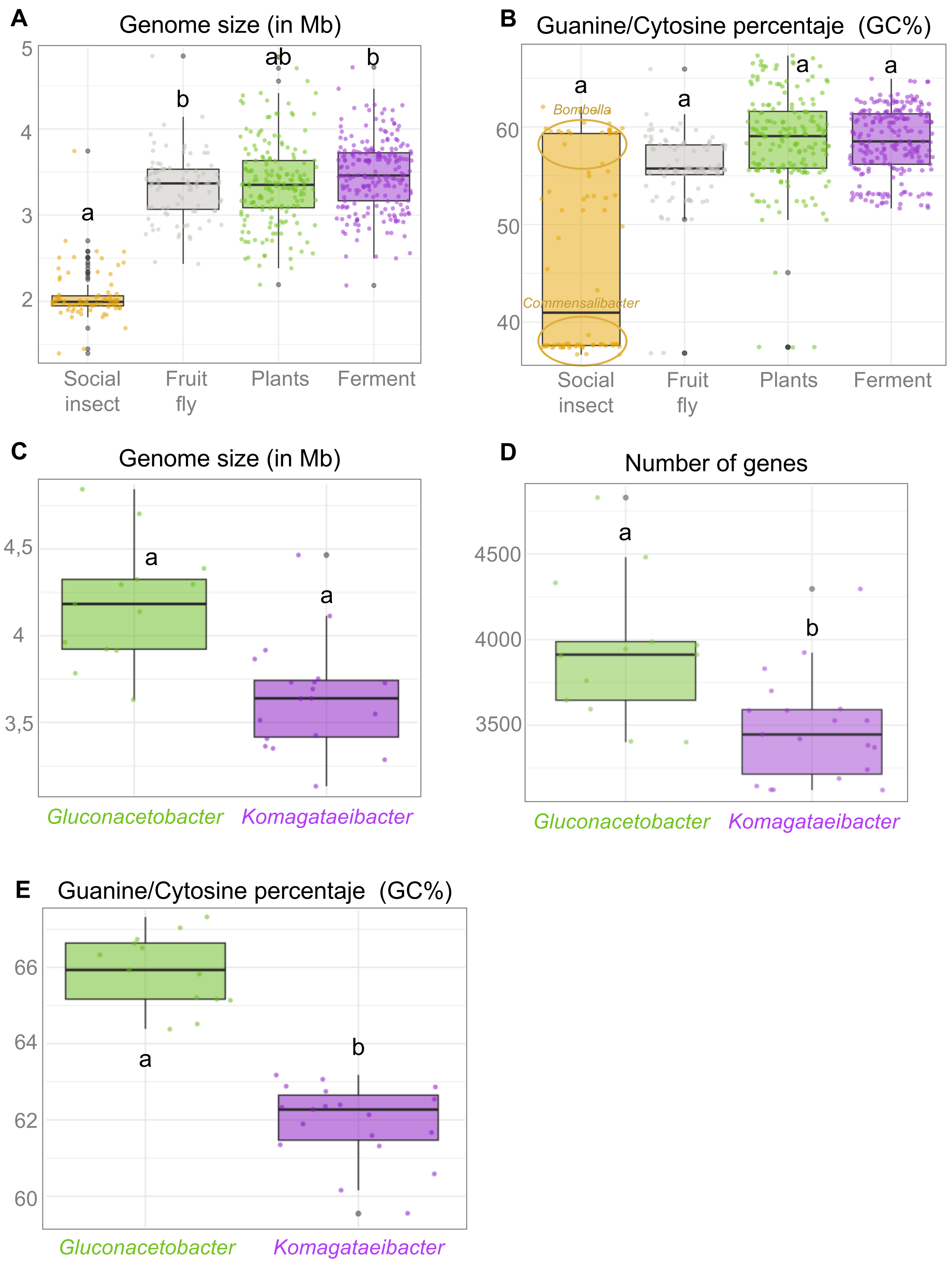

### Figure S4. Carbohydrate active enzymes (CAZyme) potential of AAB across the phylogeny, related to Figure 3.

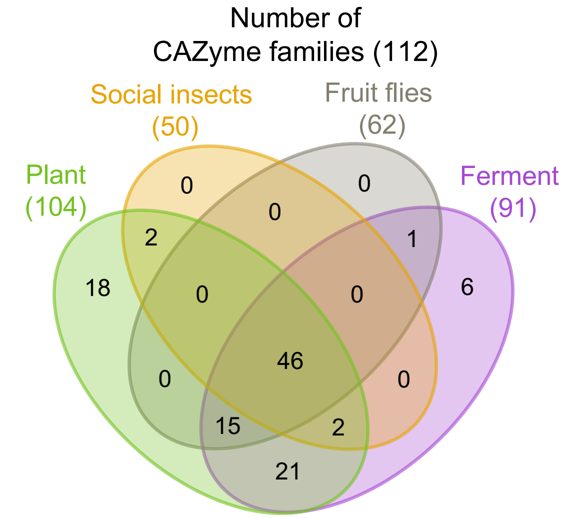
