## Supplementary material for "Evolutionary genomics reveals plant-associated origins of acetic acid bacteria in fermented food": Table S1. Pairwise comparisons of genomic characteristics of AAB genomes from Social insects, Fruit fly, Plant and Ferments, related to Figure 2.

**A**

Genome size
pairwise comparisons

| Comparison | estimate | t.ratio | p.adj |
|---|---|---|---|
| Ferment-Fly | -0,0400 | -1,438 | 0,4784 |
| Ferment-Insect | 0,1671 | 3,041 | 0,0150 |
| Ferment-Plant | 0,0335 | 1,709 | 0,3231 |
| Fly-Insect | 0,2071 | 3,721 | 0,0017 |
| Fly-Plant | 0,0735 | 2,561 | 0,0557 |
| Insect-Plant | -0,1336 | -2,526 | 0,0607 |

**B**

GC% pairwise comparisons

| Comparison | estimate | t.ratio | p.adj |
|---|---|---|---|
| Ferment-Fly | 0,0000 | 0,001 | 1,0000 |
| Ferment-Insect | 0,0008 | 0,565 | 0,9423 |
| Ferment-Plant | 0,0005 | 0,998 | 0,7508 |
| Fly-Insect | 0,0008 | 0,541 | 0,9489 |
| Fly-Plant | 0,0005 | 0,716 | 0,8906 |
| Insect-Plant | -0,0003 | -0,199 | 0,9972 |

**C**

Number of pseudogenes
pairwise comparisons

| Comparison | estimate | t.ratio | p.adj |
|---|---|---|---|
| Ferment-Fly | -0,106 | -0,361 | 0,9838 |
| Ferment-Insect | 2,159 | 7,442 | <0,0001 |
| Ferment-Plant | 0,359 | 2,069 | 0,1703 |
| Fly-Insect | 2,265 | 6,322 | <0,0001 |
| Fly-Plant | 0,465 | 1,667 | 0,3463 |
| Insect-Plant | -1,800 | -6,761 | <0,0001 |

**D**

Total number of genes
pairwise comparisons

| Comparison | estimate | t.ratio | p.adj |
|---|---|---|---|
| Ferment-Fly | -0,0304 | -0,844 | 0,8333 |
| Ferment-Insect | 0,3055 | 3,883 | 0,0010 |
| Ferment-Plant | 0,0502 | 2,030 | 0,1825 |
| Fly-Insect | 0,3360 | 4,340 | 0,0002 |
| Fly-Plant | 0,0806 | 2,205 | 0,1277 |
| Insect-Plant | -0,2554 | -3,291 | 0,0070 |
