## Supplementary material for "Evolutionary genomics reveals plant-associated origins of acetic acid bacteria in fermented food": Table S2. Carbohydrate active enzymes (CAZyme) potential of AAB across from Social insects, Fruit fly, Plant and Ferments, related to Figure 3.

**A**

Number of CAZymes pairwise comparisons

| Comparison | estimate | t.ratio | p.adj |
|---|---|---|---|
| Ferment-Fly | 0,0032 | 0,095 | 0,9997 |
| Ferment-Insect | 0,1592 | 2,447 | 0,0736 |
| Ferment-Plant | -0,0022 | -0,094 | 0,9997 |
| Fly-Insect | 0,1560 | 2,371 | 0,0880 |
| Fly-Plant | -0,0054 | -0,157 | 0,9986 |
| Insect-Plant | -0,1614 | -2,578 | 0,0534 |

**B**

Adonis test for pairwise comparisons of Beta diversity

| Comparison | df | Sum quares | F.Model | R2 | p-value | p.adjusted |
|---|---|---|---|---|---|---|
| Ferment vs Plant | 1 | 0,1463 | 3,1677 | 0,0307 | 0,0050 | 0,0300 |
| Ferment vs Fly | 1 | 0,0558 | 1,3696 | 0,0283 | 0,1486 | 1,0000 |
| Ferment vs Insect | 1 | 0,9720 | 24,6530 | 0,2912 | 0,0010 | 0,0060 |
| Plant vs Fly | 1 | 0,0932 | 1,9432 | 0,0282 | 0,0520 | 0,2167 |
| Plant vs Insect | 1 | 1,0216 | 22,2987 | 0,2180 | 0,0010 | 0,0060 |
| Fly vs Insect | 1 | 0,2800 | 7,8742 | 0,2258 | 0,0010 | 0,0060 |
